# Motor sequence analysis as a sensitive biomarker of dopaminergic degeneration in a non-human primate model of parkinsonism

**DOI:** 10.64898/2026.04.23.720333

**Authors:** Laís Resque Russo Pedrosa, Leon Claudio Pinheiro Leal, José Augusto P. C. Muniz, Arthur Gonsales da Silva, Deiweson Souza-Monteiro, Rafael Rodrigues Lima, Bruno D. Gomes, Lane V. Krejcová

**Author notes:** These authors contributed equally (Laís Resque Russo Pedrosa, Leon Claudio Pinheiro Leal). **Correponding author: Name**: Bruno Duarte Gomes, **Address**: Avenida Perimetral, 2-224, Room 238, Guamá, Belém – PA, Brazil, 66077-830, **Mail**.

## Abstract

Parkinson’s disease is characterized by progressive dopaminergic degeneration, yet motor symptoms emerge only after substantial neuronal loss - a dissociation that challenges the sensitivity of conventional behavioral endpoints in preclinical models. Here, we present a proof-of-principle study establishing a graded hemiparkinsonism model in adult male capuchin monkeys (*Sapajus apella*) through unilateral, MRI-guided stereotaxic injection of 6-hydroxydopamine into the substantia nigra pars compacta. Three toxin concentrations (4, 10, and 40 mg/mL; n = 3) were tested alongside a vehicle-injected sham control (n = 1). Motor function was assessed longitudinally before and after surgery using a three-task battery comprising the Staircase test, Tube test, and Brinkman board, capturing complementary dimensions of motor functions, including gross lateralization, forelimb use asymmetry, and fine digit coordination. Critically, we introduce a novel sequence-deviation metric applied to Brinkman board performance data to quantify disruption in the spatial organization of pellet retrieval independently of task success. Post-surgical tyrosine hydroxylase immunohistochemistry combined with optical fractionator stereology revealed ipsilateral dopaminergic cell losses of 47%, 59%, and 44% relative to the contralateral hemisphere across the three treated animals, with the sham showing no meaningful hemispheric difference. Behavioral impairments were heterogeneous and strategy-dependent: task completion rates were largely preserved, whereas fine motor strategy analysis revealed post-lesion increases in retrieval sequence disorganization in two of three animals. Exploratory regression analyses suggested that strategy-level metrics were more sensitive to nigrostriatal degeneration than global performance measures. These findings demonstrate that capuchin monkeys subjected to unilateral 6-hydroxydopamine lesions reproduce clinically relevant features of hemiparkinsonism and that motor sequence analysis constitutes a sensitive readout of subclinical dopaminergic dysfunction, and can outperform conventional performance-based metrics detecting early motor alterations, therefore a potential biomarker of subclinical dopaminergic dysfunction, with implications for early detection paradigms in Parkinson’s disease research.

## 1. Introduction

Parkinson’s disease (PD) is the second most prevalent neurodegenerative disorder globally, yet its defining motor manifestations emerge only after the nigrostriatal system has sustained substantial, largely irreversible damage. Approximately 50–60% of dopaminergic neurons in the substantia nigra pars compacta and up to 80–85% of striatal nerve terminals are lost before overt motor dysfunction becomes clinically apparent - a threshold that reflects the remarkable compensatory plasticity within the nigrostriatal system (Braak et al., 2003; Zigmond et al., 1990). Consequently, behavioral measures based solely on task success or completion rate may systematically underestimate the severity and organization of underlying motor dysfunction, precisely because preserved performance can mask substantial reorganization of motor control strategies.

Rodent models provide valuable mechanism insights for understanding the neural basis of PD, but the behavioral repertoire is limited for translating many clinically relevant outcome measures (Prasad & Hung, 2020).Fine motor assessments, upper limb dexterity tasks analogous to human clinical evaluations, and many clinically relevant motor and non-motor PD features cannot be adequately modeled in rodents with the same validity achievable in Non-Human Primates (NHPs) (Bezard et al., 2025). Among NHP species, capuchin monkeys (*Sapajus apella*) present a particularly compelling model for fine motor investigation. These animals possess a highly organized motor cortex with well-defined digit representations, exhibit spontaneous tool-use behavior in natural conditions, and have dopaminergic neurons containing neuromelanin, a feature shared with humans and associated with selective vulnerability to oxidative stress (Pedrosa et al., 2024). These features support the implementation of high-resolution fine motor tasks with a high degree of translational relevance.

6-Hydroxydopamine (6-OHDA) was the first neurotoxin used to model Parkinson’s disease experimentally and remains one of the most mechanistically characterized tools for inducing selective dopaminergic loss. Unlike MPTP, which can be administered systemically and exploits species-specific monoamine oxidase B activity, 6-OHDA does not cross the blood–brain barrier and therefore requires direct stereotaxic injection into dopaminergic target structures (Emborg, 2007). When delivered to the substantia nigra (SN), 6-OHDA is selectively taken up via the dopamine transporter and triggers retrograde nigrostriatal degeneration through oxidative stress and mitochondrial complex I inhibition. Unilateral injection produces an asymmetric hemiparkinsonism (HP) phenotype in which the contralateral hemisphere remains intact. This provides a within-subject internal control for both behavioral and histological comparisons, an experimental advantage that bilateral models and systemic neurotoxins cannot offer. The ability to titrate lesion severity through concentration adjustments further enables investigation of the relationship between graded dopaminergic loss and motor impairment across a continuum that spans sub-threshold, threshold, and suprathreshold degeneration.

Motor behavior is inherently multidimensional, encompassing not only the ability to complete the task successfully but also the way the task has been completed - coordination, sequencing, and spatial organization of movements (Schmidlin et al., 2011). Dopaminergic degeneration may affect these distinct components differently, hence neither reductions in task success nor speed fully capture disruptions in motor planning and control. A comprehensive characterization of motor function, therefore, requires integrating complementary behavioral dimensions, rather than relying on a single endpoint measure. In this study, we operationalized this multidimensional framework using a battery of three complementary motor tasks - the Staircase test, the Tube test, and the Brinkman board - each capturing a distinct behavioral dimension ranging from gross forelimb lateralization to fine digit coordination and motor sequence organization. Critically, a novel sequence-deviation metric specifically designed to capture disruptions in motor sequence organization independently of task success was applied to Brinkman board data to quantify disruption in the spatial organization of pellet retrieval, independently of total task performance.

Given the translational motor phenotype fidelity of capuchin monkeys and the mechanistic tractability of focal 6-OHDA lesions, we aimed to establish a graded HP model in adult male *Sapajus apella* using MRI-guided stereotaxic 6-OHDA injections at three concentrations (4, 10, and 40 mg/mL) into the SN. This proof-of-principle study combined longitudinal behavioral assessments across three complementary motor tasks with Tyrosine Hydroxylase (TH) immunohistochemistry and stereological analysis through the optical fractionator technique to quantify dopaminergic neurons in the SN. By integrating measures of limb asymmetry, task latency, grasping strategy, and motor sequence organization, we aimed to characterize how distinct dimensions of motor performance are differentially affected by graded nigrostriatal degeneration, and test the hypothesis that strategy-level metrics are more sensitive than conventional performance measures for detecting dopaminergic dysfunction.

## 2. Methods

### 2.1 Animals

Four adult male capuchin monkeys (*Sapajus apella*) were obtained from the National Primate Center (Centro Nacional de Primatas, IEC) Ananindeua, Pará, Brazil. The animals had a mean age of 19.66 ± 5.75 years and a mean body weight of 4.30 ± 0.99 kg. All animals were clinically healthy with no prior history of neurological disease. They were single-housed in standard cages (2.5 × 2.0 × 2.5 m) under a 12 h light/dark cycle. Their diet included laboratory chow specifically formulated for non-human primates, supplemented with fresh fruits and natural juice. Water was available *ad libitum*. Environmental enrichment was provided in accordance with the International Guidelines for non-human primate’s welfare.

All experimental procedures were conducted in accordance with Directive 2010/63/EU of the European Union and ARRIVE guidelines and were approved by the Ethics Committee for the Use of Animals at the Evandro Chagas Institute (CEUA/IEC), under protocol numbers 45/2016 and 37/2018.

### 2.2 Behavioral Assessment

Animals underwent behavioral assays to evaluate bilateral motor function, including both fine and gross motor coordination, and side dominance. Subjects were tested individually three times per week, with sessions scheduled alternately and always conducted prior to daily feeding. Behavioral assessments were conducted both before (baseline) and after surgery to analyze changes in motor performance. The three-task battery was the Staircase, Tube, and Brinkman board tasks. All behavioral assessments were performed by the same experimenter in order to minimize inter-rater variability.

#### 2.2.1 Staircase test

An adapted staircase apparatus for non-human primates was used (J. Marshall et al., 2002; J. W. Marshall & Ridley, 2003). It corresponds to the Valley version of the apparatus, in which a central opening for accessing the reward is placed on ascending steps toward the sides. A total of 120 sessions were conducted across pre- and post-surgical phases, each limited to a maximum duration of 3 minutes for reward retrieval. The following parameters were scored for each side: first-retrieval latency, time to retrieve all rewards, number of cross-hand reaches (using the opposite hand for a given side), and number of dropped rewards.

#### 2.2.2 Tube test

The tube test was performed using a PVC tube with two upper openings, filled with an edible reward of creamy texture (e.g. peanut butter). Each animal was given 1.5 minutes per session to retrieve the reward using either forelimb. Twenty-five sessions were conducted before and after surgery. The following parameters were assessed: time spent using the dominant and non-dominant hand, number of grasps for each hand, frequency of index finger use, and frequency of use of other fingers.

#### 2.2.3 Brinkmann board test

The modified Brinkman board was used to quantitatively assess fine motor coordination by measuring the ability to retrieve small pellets (Rouiller et al., 1998; Yamanaka et al., 2021). The apparatus consists of an acrylic board (22 x 12 cm) with 50 rectangular wells —25 arranged vertically and the remaining 25 horizontally—each filled with a flavored pellet.

The board was positioned at a 45° angle on a wooden table in front of the animal, which was isolated in the upper compartment of its cage without unilateral restriction. Non-human primates were trained to retrieve using a precise pincer grip (thumb opposed to index finger). Twenty-five sessions were conducted in both the pre- and post-surgical periods. No time limit was imposed for completing the test. Motor performance was evaluated by the number of pellets collected per session using the dominant and non-dominant hands, as well as through a quantitative index of motor sequence organization adapted from a previous work (Schmidlin et al., 2011).

### 2.3 Lesions

Animals were initially anesthetized with an intramuscular injection of xylazine (100 mg/Kg) and ketamine (100 mg/Kg). After animal positioning in the stereotaxic frame, anesthesia was maintained with an intravenous infusion of the same agents under constant monitoring by the veterinary staff.

MRI-guided stereotaxic targeting was performed individually for each subject based on anatomical landmarks and behavioral lateralization, targeting the dominant side identified during baseline behavioral testing. Subsequent 6-OHDA injections were executed using subject-specific MRI data, following the methodology and stereotaxic coordinates described by Pedrosa et al., 2024.

Animals in the HP group received unilateral 6-OHDA injections dissolved in 0.01% ascorbic acid in saline. A volume of 2 µL was delivered at each of four equidistant sites within the SN (injection rate: 0.5 µL/min). The SHAM animal received vehicle-only injections (0.01% ascorbic acid in saline) at the same stereotaxic coordinates. The 6-OHDA concentrations administered to each animal are provided in Table 1.

**TABLE 1.** The HP animals received different concentrations of 6-OHDA (4, 10, and 40 mg/mL) to evaluate potential motor impairments. Lesions were induced in the contralateral hemisphere to the preferred hand assessed previously. The SHAM animal received vehicle injections only (0.01% ascorbic acid).

| Subject | Condition | 6-OHDA<br>Concentration<br>(mg/ml) | Lesioned<br>side |
| --- | --- | --- | --- |
| AM-BEN | HP | 4 | Left |
| AM-BEG | HP | 10 | Right |
| AM-AOR | HP | 40 | Left |
| AM-AQX | SHAM | - | Left |

### 2.4 Tyrosine Hydroxylase (TH) Immunohistochemistry

After the completion of the behavior experiments, the animals were euthanized by anesthetic overdose using a combination of ketamine hydrochloride (100 mg/kg) and xylazine hydrochloride (50 mg/kg). To fix the nervous tissue, the subjects were transcardially perfused with 0.9% saline, followed by 4% paraformaldehyde (PFA). Brains were extracted, post-fixed, and coronally sectioned (50µm) with a vibratome.

For TH immunohistochemistry, the sections were pre-treated for antigen retrieval using 0.3% boric acid solution and washed three times in PBS for 5 minutes each. Subsequently, they were incubated for 20 minutes in 10% serum for blocking. The sections were then incubated in a solution containing anti-tyrosine hydroxylase primary antibody (1:1000 in PBS, pH 7.2-7.4) for 72 hours (Anti-TH (Ab-5) Rabbit pA, PC38-100ul – Santa Cruz Biotechnology, Inc.).

After incubation, the sections were removed from the primary antibody solution and washed four times in 0.1M phosphate buffer for 5 minutes each. They were then incubated overnight with the secondary antibody. The next day, the sections were washed three times in 0.1M PBS for 5 minutes each, followed by blocking of endogenous peroxidase using 0.3% hydrogen peroxide (H₂O₂) in 0.1M phosphate buffer for 10 minutes. After three more washes, the sections were incubated in an avidin-biotin enzyme complex solution (ABC kit, PK-4000, Vector, Burlingame, CA) for 1 hour. Then, they were rewashed and subjected to the peroxidase detection reaction using DAB (diaminobenzidine – Sigma-Aldrich, Inc) as the chromogen. Finally, the sections were washed in a 0.1M phosphate buffer, dehydrated, cleared, and mounted.

### 2.5 Microscopic Analysis and Cell Counting Techniques

Dopaminergic neurons in the SN were quantified using the optical fractionator method, a precise stereological approach for quantifying cell populations that combines the functionality of an optical dissector and a fractionator (Bonthius et al., 2004; M. J. West, 1993, 1999). One of the main advantages of this method is its resistance to histological alterations such as tissue shrinkage or expansion induced by lesions (M. West et al., 1991).

In the histological sections, we precisely identified the layers of SN, positioning counting probes and capturing digital images. For this, we used a low-magnification objective lens (4×) on a NIKON Eclipse 80i microscope (Nikon, Tokyo, Japan) equipped with a motorized stage (MAC200, Ludl Electronic Products, Hawthorne, NY, USA). The system was connected to a computer running StereoInvestigator software (MicroBrightField, Williston, VT, USA, https://www.mbfbioscience.com/products/stereo-investigator, accessed on February 15, 2025), allowing the precise digital recording and analysis of the x, y, and z coordinates of the selected points.

To ensure accurate neuron identification with the dissector probe, we replaced the low-magnification objective with a high-resolution oil immersion objective (100×, Nikon, NA 1.3, DF = 0.19 µm). This adjustment enabled unequivocal neuron cell counting.

Moreover, the thickness of each section was carefully measured at each counting site using the high-resolution objective, allowing precise delimitation of the upper and lower planes. Given the variability in thickness and cell distribution across sections, the total number of cells of interest was adjusted based on that thickness. Only neuronal cell bodies clearly visible within the counting frame or crossing the acceptance line without touching the rejection line were included, following previously established methodological criteria (Gundersen & Jensen, 1987). To ensure comprehensive and unbiased sampling, the counting frames were distributed systematically and randomly within a grid.

All sections from the first appearance to the decussation of the SN were used, generating between 6 and 8 sections with intervals of 6 for each technique and each animal. Depending on the technique, the counting box size ranged from 50 to 150 µm², with 15 to 20 boxes per ROI (region of interest). The optical dissector was adjusted based on the actual section thickness, which could vary depending on the shrinkage produced by each technique. A Gundersen coefficient error < 0.07 was adopted. Counting boxes were consistently positioned within a grid in a randomized yet systematic manner to ensure comprehensive coverage and unbiased sampling.

Percent cell loss was computed in percent loss relative to the contralateral side:

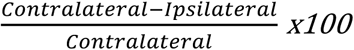

### 2.6 Behavioral statistical analysis

#### 2.6.1 Staircase and Tube test analysis

Data were tested for normality using the D’Agostino–Pearson test. Non-normally distributed data were analyzed with the Wilcoxon signed-rank test for within-subject comparisons and with the Kruskal–Wallis test, followed by Dunn’s post hoc test with Bonferroni correction, for inter-group comparisons. Data are reported as median and interquartile range.

#### 2.6.2 Brinkman sequence analysis

These data were visualized as heatmaps to assess the consistency of the retrieval strategy across sessions and to evaluate its alteration following the intervention. The retrieval order was represented as a session-retrieval-step matrix. It contains the pellet number retrieved at the corresponding step.

To quantify sequence disorganization, a deviation score was computed for each retrieval event as the difference between the retrieved pellet number and the ordinal retrieval step. Therefore, the value of zero indicated perfect agreement with the canonical sequence, whereas nonzero values reflected deviations from that order.

For each session, the deviation variance metric was calculated across all retrieval events and used as sequence variability, with higher values indicating greater sequence instability. Session-level deviation variance values were compared between pre- and post-intervention conditions for each animal. Statistical comparisons were performed using the Mann–Whitney U test.

### 2.6.3 Behavioral Data Processing and Strategic Analysis

Behavioral performance across tasks (Staircase success rate, Tube test asymmetry, Brinkman board variability) was normalized to each animal’s baseline. Functional impairment was calculated as a percentage change from baseline:

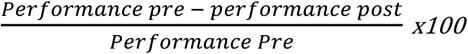

Negative values indicate animals that maintained or exceeded baseline performance, potentially reflecting motor adaptation or task-related learning.

To quantify alterations in motor organization, a variability metric was derived from the Brinkman task. For each trial, spatial deviation was defined as the difference between the observed retrieval sequence and the canonical board order (positions 1–50). Absolute deviations were computed to avoid directional cancellation, and the mean absolute deviation was calculated per session. These values were normalized to baseline using the same formula, providing an index of post-lesional increases in sequence variability.

### 2.7 Correlation with Dopaminergic Degeneration

Dopaminergic degeneration was quantified as the percentage loss of tyrosine hydroxylase (TH)-positive neurons in the SN, based on comparisons between ipsilateral (lesioned) and contralateral (not-lesioned) hemispheres.

Animal-level behavioral changes were compared with TH loss using simple linear regression. Given the small sample size, analyses were considered exploratory, with emphasis on effect sizes and consistency across measures. The coefficient of determination (R²) was used to describe the variance explained. An animal-level heatmap was generated to summarize behavioral and histological measures.

A schematic representation of the experimental timeline, including behavioral assessment and surgical intervention, is shown in Figure 1.

**FIGURE 1.**
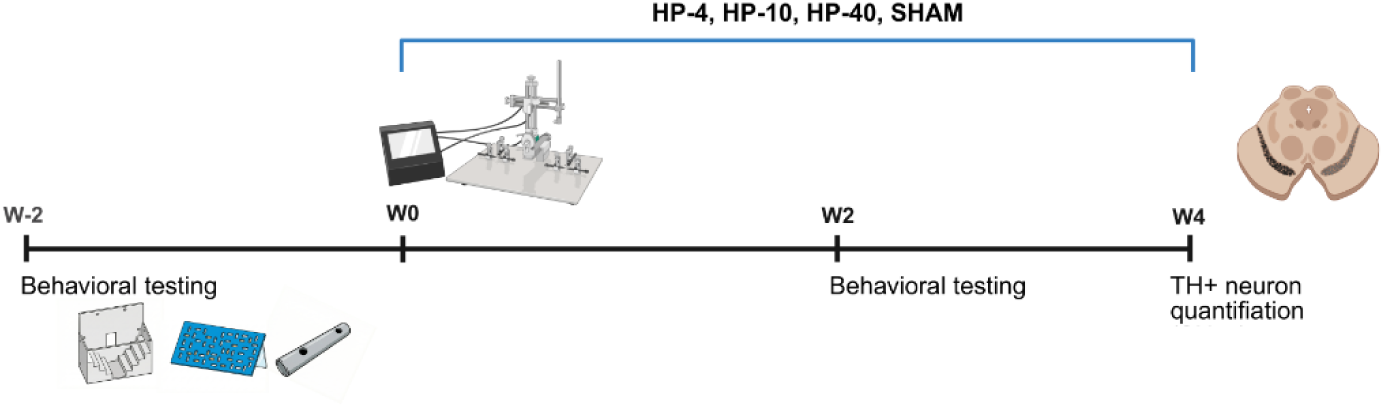
Experimental timeline for the hemiparkinsonism induction model in *Sapajus apella*. Baseline motor assessments (Staircase, Tube, and Brinkman board tests) were conducted during the pre-surgical period (approximately 2 weeks before surgery; not drawn to scale). At week 0 (W0), animals underwent MRI-guided stereotaxic surgery for unilateral 6-OHDA injection (2 µL at four equidistant SN sites; dissolved in 0.01% ascorbic acid saline) or vehicle-only injection (SHAM). Post-surgical motor assessment began at week 2 (W2). Animals were euthanized at week 4 (W4) for TH immunohistochemistry and optical fractionator stereology. Subjects: HP-4 (6-OHDA 4 mg/mL), HP-10 (6-OHDA 10 mg/mL), HP-40 (6-OHDA 40 mg/mL), SHAM (vehicle).

## 3. Results

### 3.1 Behavioral adaptations reveal preserved performance despite underlying motor deficits

To determine whether graded dopaminergic lesions induce detectable motor deficits, animals were assessed using three-task battery before and after 6-OHDA administration.

In the Staircase test, subjects were required to retrieve ten single-placed rewards. All the animals maintained their consistency in performing the task independently of the surgical condition, which indicates that overall task performance was largely preserved across animals (see Supplementary Figure 1A). However, strategy-level analysis revealed asymmetric motor adaptations, including increased reliance on the non-dominant limb and altered retrieval latency patterns following lesion (Figures 2 and 3): BEN HP 4 exhibited a compensatory behavior toward the non-dominant side, characterized by increased first-retrieval latency on the dominant side, decreased latency on the non-dominant side, and a significant post-surgical switch in hand preference. In contrast, BEG HP10 did not adopt a comparable compensatory strategy, despite an increased loss of rewards on the dominant side after surgery. Nonetheless, AOR HP 40 showed clear non-dominant compensation by a significant reduction in dominant-side performance.

**FIGURE 2.**
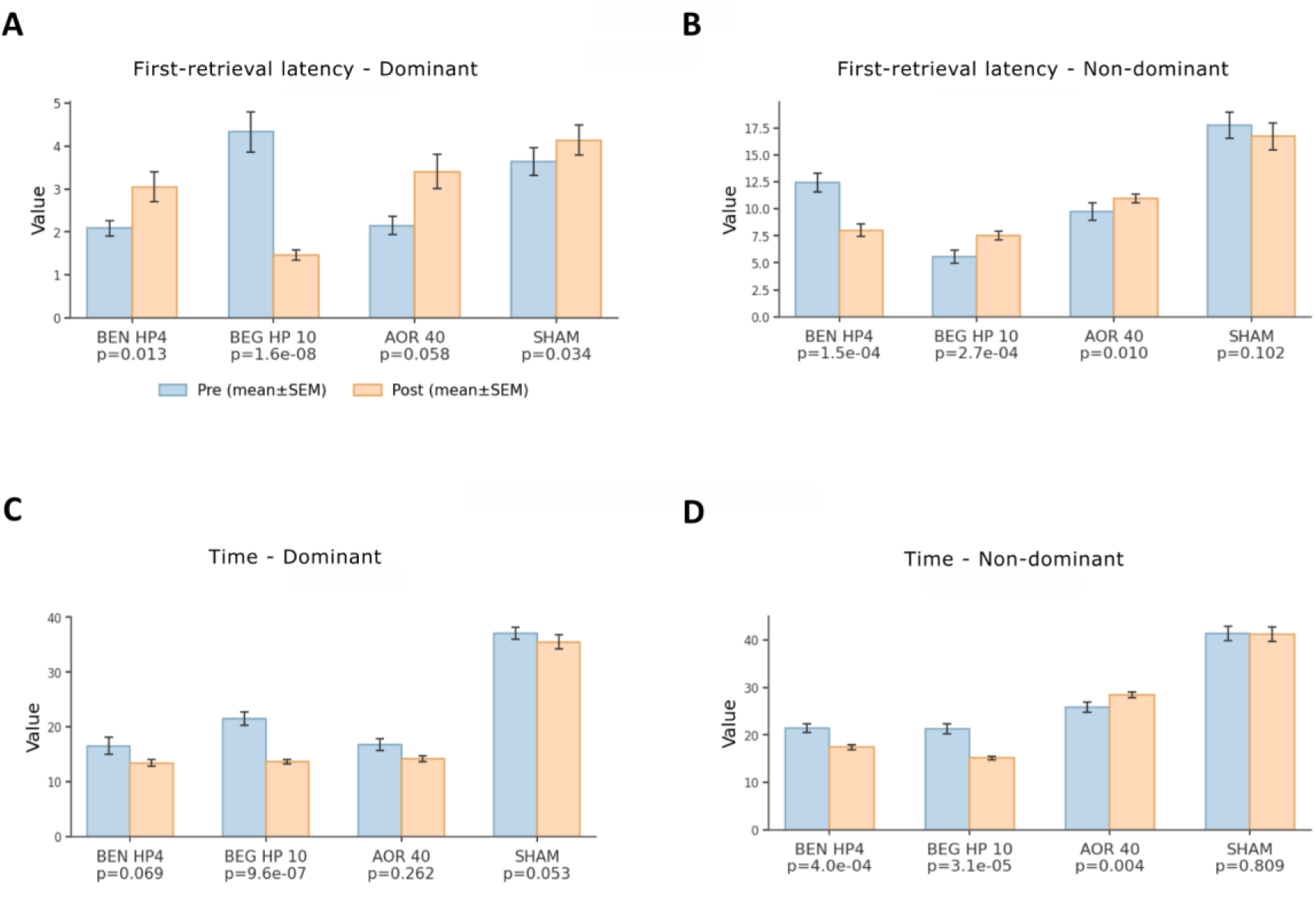
Staircase test: reaction latency and retrieval time. (A) First-retrieval latency on the dominant side. (B) First-retrieval latency on the non-dominant side. (C) Total retrieval time per session on the dominant side. (D) Total retrieval time per session on the non-dominant side. Blue bars: pre-surgical baseline; orange bars: post-surgical assessment. Data are presented as mean ± SEM (seconds). P-values from Wilcoxon signed-rank tests (pre vs. post within-subject) are shown below each subject label. N = 120 sessions per phase.

**FIGURE 3.**
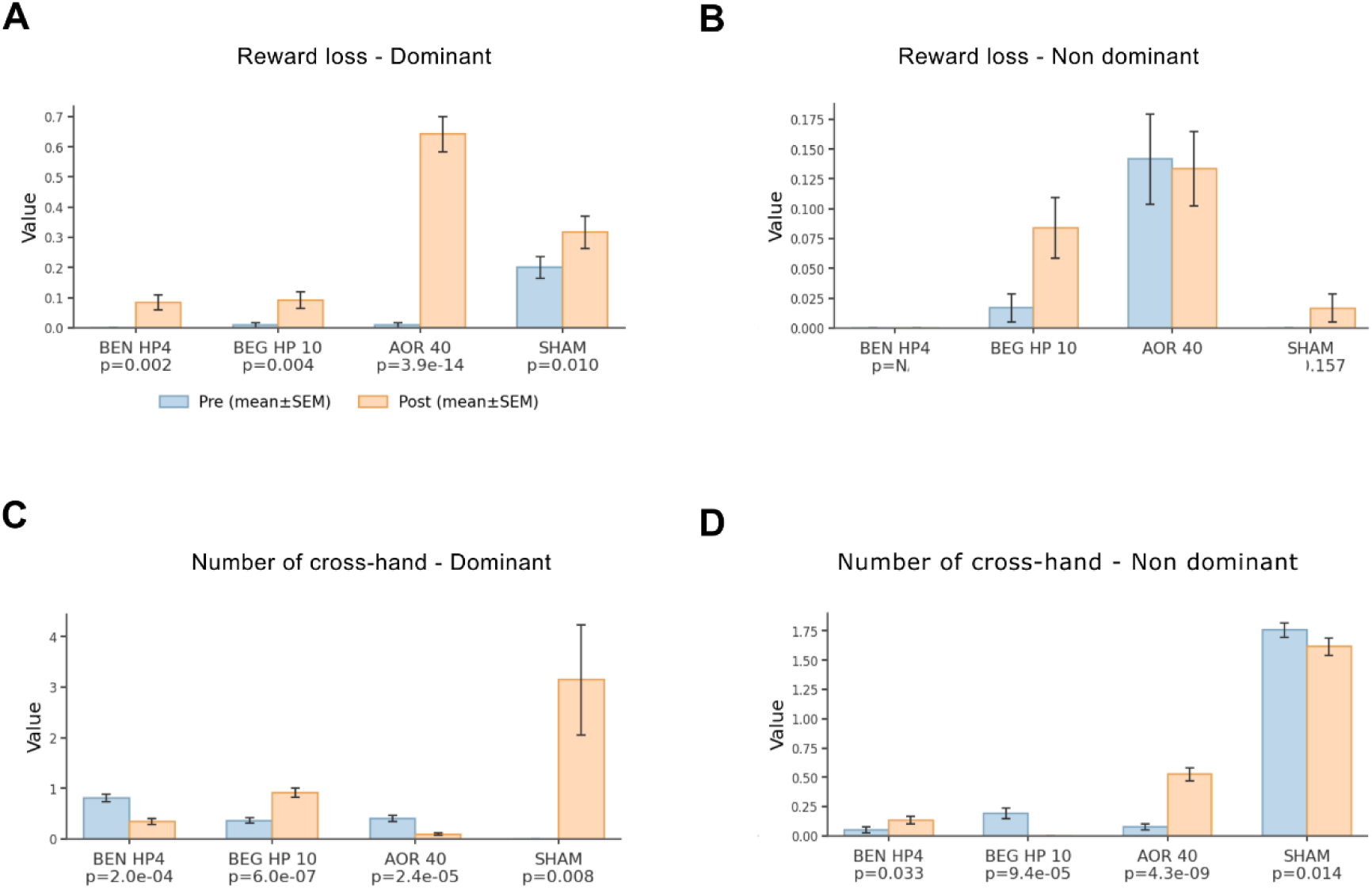
Staircase test: reward loss and cross-hand reaching. (A) Number of dropped rewards on the dominant side. (B) Number of dropped rewards on the non-dominant side. (C) Number of cross-hand reaches on the dominant side. (D) Number of cross-hand reaches on the non-dominant side. Blue bars: pre-surgical baseline; orange bars: post-surgical assessment. Data are mean ± SEM. P-values from Wilcoxon signed-rank tests are indicated below each subject label. Note: the SHAM animal exhibited a significant post-surgical increase in dominant-side cross-hand reaches (panel C, p=0.008), which may reflect a non-specific effect of the surgical procedure independent of dopaminergic lesion.

The Tube test probes motor execution under a limited time per session. In this task, animals exhibited post-lesion shifts in forelimb use, characterized by reduced engagement of the dominant limb and compensatory recruitment of the non-dominant side (Figure 4): BEN HP 4 and BEG HP 10 exhibited such post-surgical strategies, with reduced time spent and fewer grips on the dominant side, concomitant with increased use of the non-dominant side. However, AOR HP40 increased grip counts on both sides but exhibited a post-surgical bias, spending more time on the dominant side.

**FIGURE 4.**
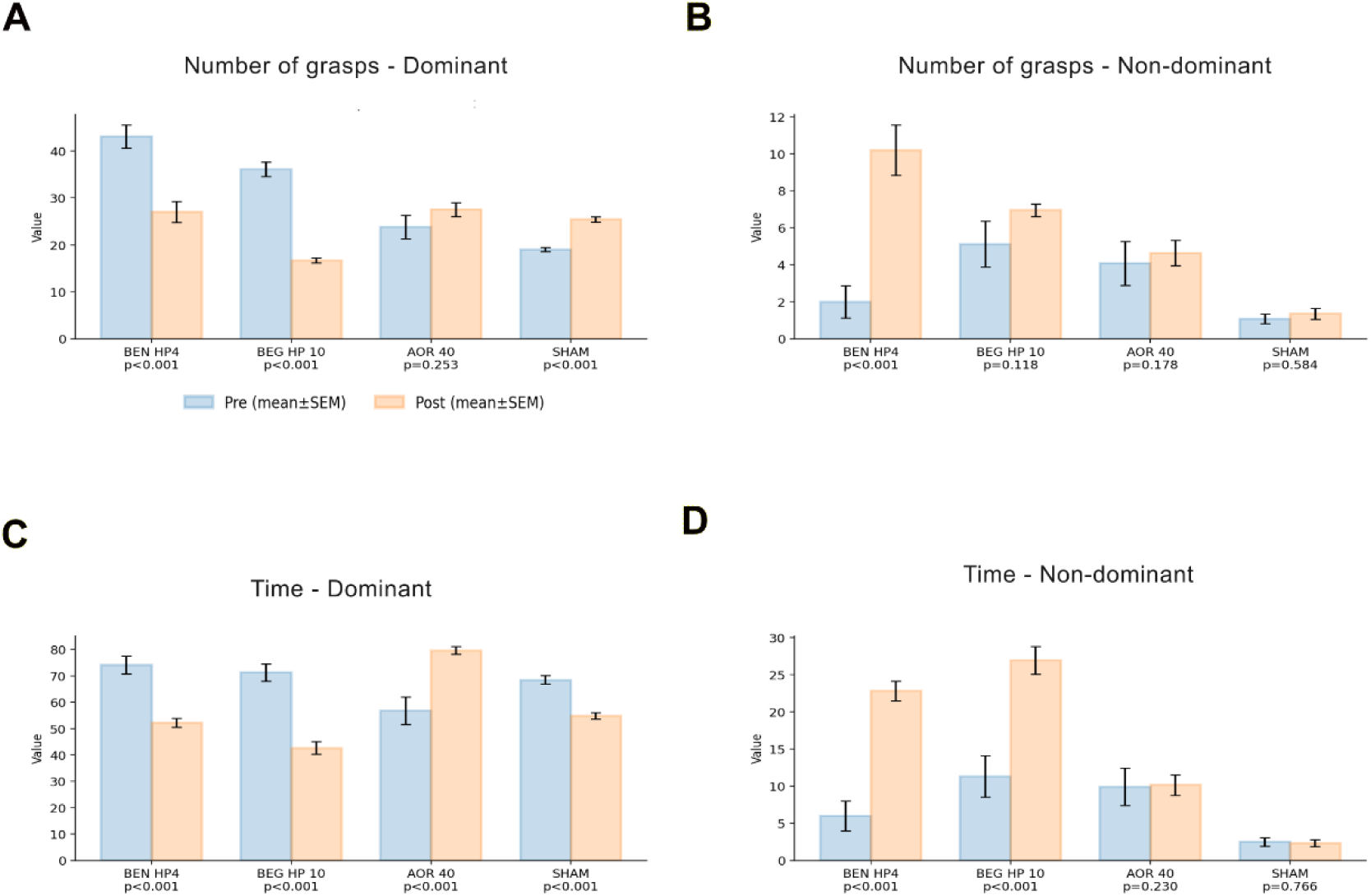
Tube test: forelimb grasping and duration. (A) Number of grasps using the dominant hand per session. (B) Number of grasps using the non-dominant hand per session. (C) Total time (seconds) spent using the dominant hand per session. (D) Total time (seconds) spent using the non-dominant hand per session. Blue bars: pre-surgical baseline; orange bars: post-surgical assessment. Data are mean ± SEM. P-values from Wilcoxon signed-rank tests are indicated below each subject label. N = 25 sessions per phase.

Collectively, these findings indicate that 6-OHDA lesions induce heterogeneous but consistent strategy-level motor adaptations, in which compensatory recruitment of the non-dominant side coexists with subtle signs of bradykinesia on the dominant side. Importantly, preserved task completion masked underlying alterations in motor strategy.

Fine motor performance was further evaluated using the Brinkman board task, with particular emphasis on motor sequence organization. For the total score in the Brinkman board task, Wilcoxon analyses revealed significant pre–post differences within the experimental groups (Supplementary Figure 1B). While overall performance showed limited changes across animals, sequence-level analysis revealed marked changes in motor organization after lesion: significant changes were observed in the BEG HP 10 and AOR HP 40 for both right and left sides, indicating a clear effect of the intervention in these groups. No significant pre–post differences were found in the BEN and SHAM groups (p > 0.05). Additionally, right–left comparisons demonstrated significant asymmetry in the post-intervention condition for the BEG HP 10 and AOR HP 40 (p < 0.05). In contrast, no significant differences between sides were observed in the BEN and SHAM (p > 0.05).

Fine motor function was assessed using the Brinkman board task, in which the sequence of pellet retrieval across 50 board positions was recorded for each session. Sessions were categorized into pre- and post-intervention phases according to the experimental design. Sequence variability differed across individuals, with some animals exhibiting increased post-lesion disorganization, while others displayed high baseline variability: BEG HP 10 showed greater variance in deviation than the other subjects, suggesting that this animal adopted a distinct retrieval strategy at baseline. Comparisons between pre- and post-intervention phases were significant for BEN HP 4 and AOR HP 40, but not for BEG HP 10 (Figure 5A).

**FIGURE 5.**
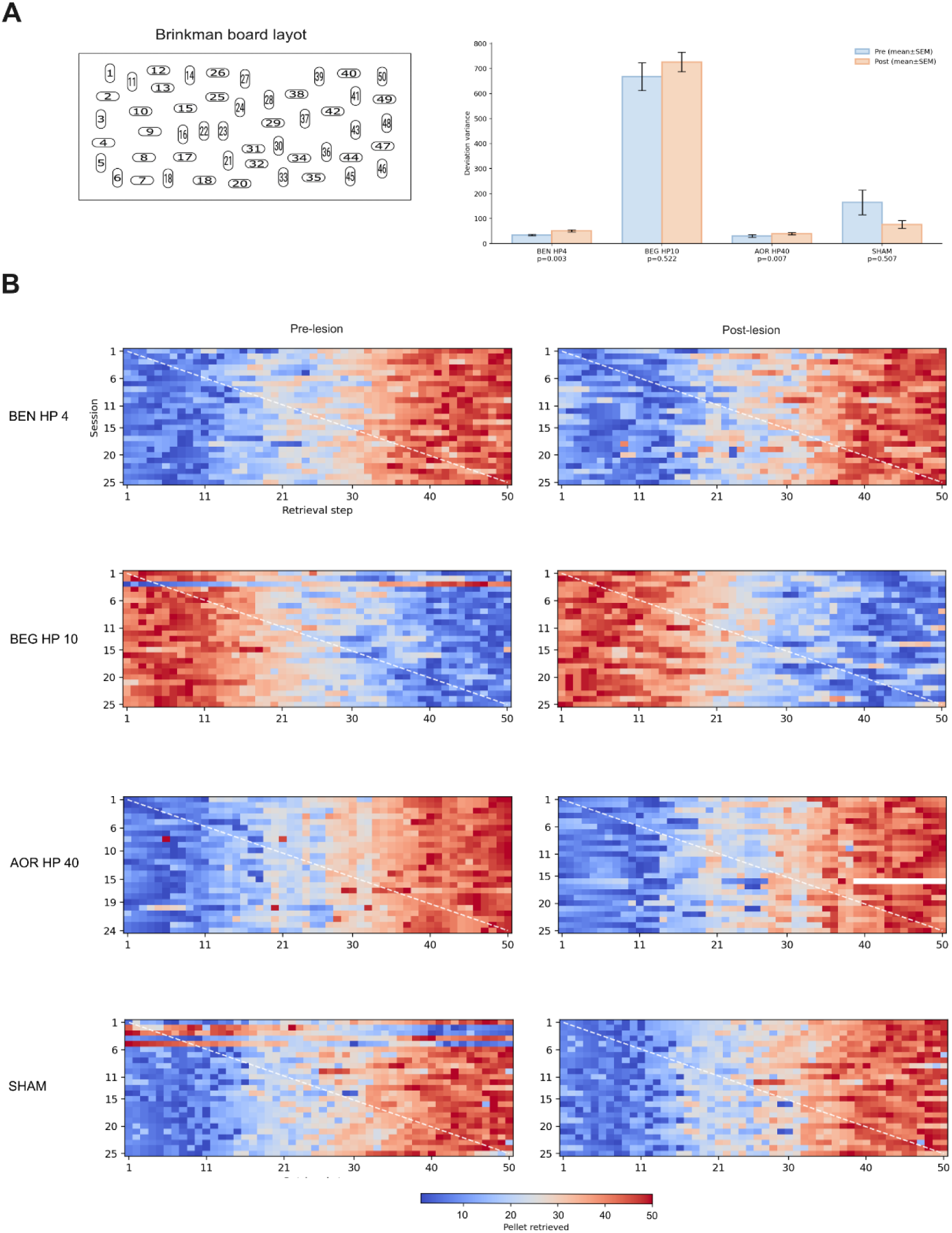
Brinkman board: motor sequence analysis. (A) Left: schematic of the 50-slot Brinkman board (22 × 12 cm acrylic board; 25 vertically and 25 horizontally oriented wells). Right: session-level deviation variance (mean ± SEM) before (blue) and after (orange) surgical intervention. Deviation variance quantifies within-session deviations between actual and expected retrieval order (see Methods 2.7); higher values indicate greater sequence disorganization. P-values from Mann-Whitney U tests (pre vs. post) are shown below each subject label. (B) Retrieval sequence heatmaps for the dominant hand across pre- (left) and post-lesion (right) sessions. Each row represents one session; each column represents an ordinal retrieval step (1–50). Color encodes the pellet number retrieved at each step (color scale: blue = low pellet number, red = high pellet number; range 10–50). The white dashed diagonal represents the ideal sequential strategy, where pellet number equals retrieval order. Organized performance appears as a smooth blue-to-red diagonal gradient; disorganization manifests as color fragmentation across the heatmap. N = 25 sessions per phase. Note: BEG HP10 showed high baseline sequence variability in both phases; white cells in AOR HP40 post-lesion indicate excluded sessions.

To visualize individual pellet retrieval strategies within each session, heatmaps were generated to represent the spatial distribution of retrieval order across board positions, relative to the default task strategy. More organized performance was characterized by smoother spatial gradients and localized clusters of similar values, indicating a consistent progression of pellet retrieval across neighboring slots.

Heatmap visualization revealed a clear disruption of spatial retrieval organization following lesion, with loss of structured sequential patterns in affected animals: In BEN HP 4 and AOR HP 40, post-lesion heatmaps became less spatially coherent, with a more irregular and fragmented distribution of retrieval order, consistent with reduced motor organization after intervention. In contrast, BEG HP 10 showed a relatively dispersed and weakly organized pattern already during the pre-intervention phase, with only minor post-intervention changes after intervention, consistent with its higher deviation variance (Figure 5B)

### 3.2 Reduction of dopaminergic neurons in 6-OHDA Hemiparkinsonian non-human primates

After behavioral assessment, TH staining was performed to evaluate the integrity of dopaminergic neurons in the SN. Using stereology and the mean section thickness estimator, we observed that unilateral 6-OHDA injections resulted in substantial and asymmetric dopaminergic cell loss across all treated animals, ranging from approximately 44% to 59% relative to the contralateral hemisphere. BEN HP 4 showed 150,022.61 TH+ cells bodies contralaterally versus 80,080.39 ipsilaterally (46.6% reduction). BEG HP 10 showed 89,326.05 contralaterally versus 36,759.49 ipsilaterally (58.8% reduction). The AOR HP 40 showed 105,697.83 contralaterally versus 59,488.58 ipsilaterally (43.7% reduction). No significant hemispheric difference was observed in the SHAM animal, confirming the specificity of the lesion (110,176.22 contralateral vs 110,334.87 ipsilateral; −0.1%), see Figure 6.

**Figure 6.**
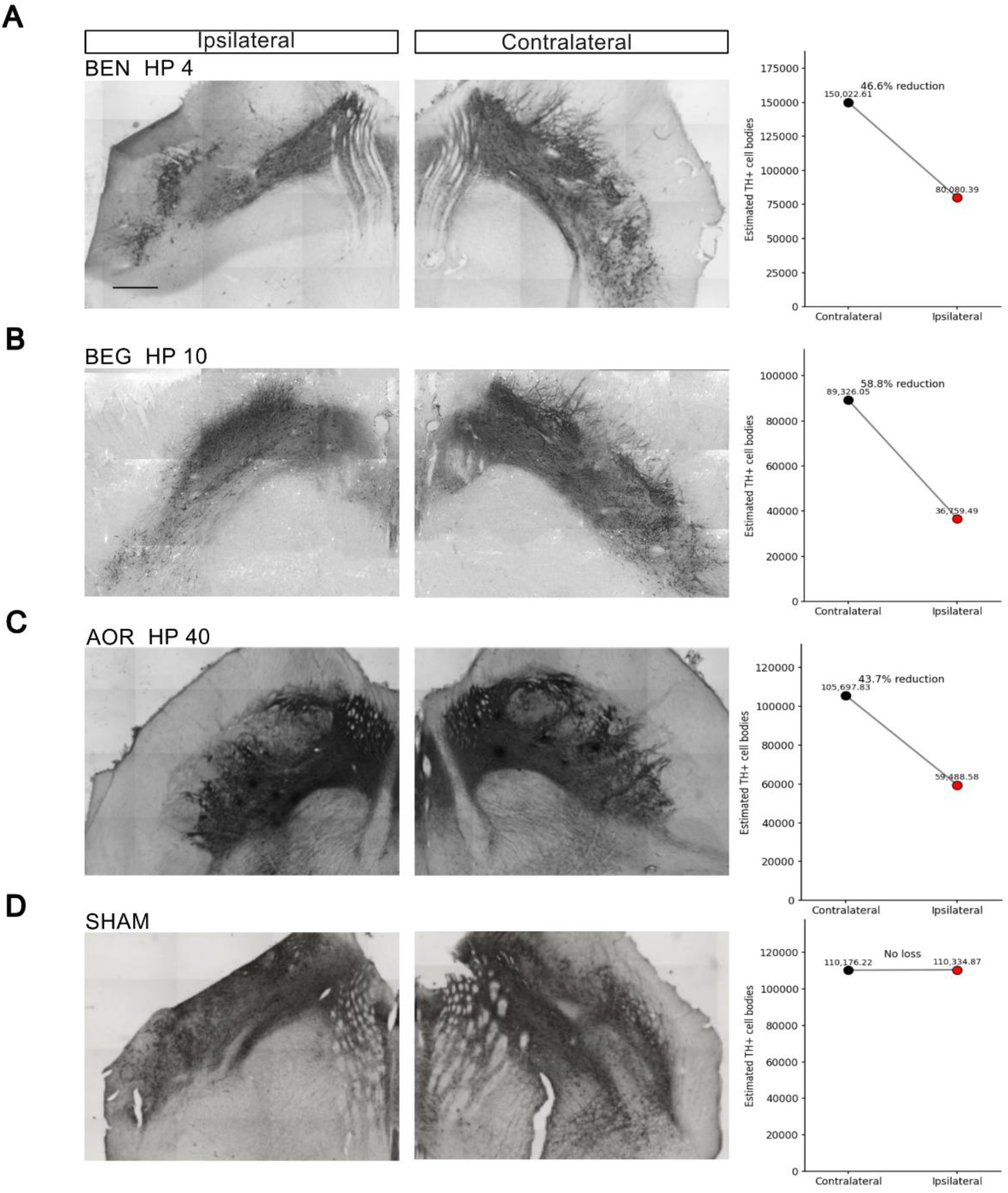
Unilateral 6-OHDA injection induces ipsilateral dopaminergic cell loss in the substantia nigra of adult *Sapajus apella*. Left: representative TH photomicrographs of the ipsilateral (lesioned) and contralateral (non-lesioned) SN from coronal vibratome sections (50 µm). Images were acquired at 2.5× magnification. Scale bar = 500 µm (shown in A; applies to all photomicrographs). Rows correspond to individual subjects: (A) BEN HP4 (6-OHDA, 4 mg/mL); (B) BEG HP10 (6-OHDA, 10 mg/mL); (C) AOR HP40 (6-OHDA, 40 mg/mL); (D) SHAM (vehicle-injected control, 0.01% ascorbic acid in saline). Right: subject-level estimates of the total number of TH+ cell bodies in the SN, obtained by the optical fractionator method with actual measured section thickness (Gundersen CE ≤ 0.07; for full stereological parameters see Methods). Black filled circles = contralateral hemisphere; red filled circles = ipsilateral hemisphere; connecting lines indicate the direction and magnitude of interhemispheric change. Percentage values indicate ipsilateral cell loss relative to the contralateral hemisphere [(contralateral − ipsilateral)/contralateral × 100]. TH, tyrosine hydroxylase; SN, substantia nigra; HP, hemiparkinsonism.

### 3.3 Exploratory association between behavioral impairment and histological lesion severity

To explore the relationship between dopaminergic degeneration and behavioral outcomes, we compared histological and motor behavior metrics across subjects (see the graphs in Figure 7). As previously mentioned, conventional performance-based metrics for motor assessment (e.g. score, time) showed limited association with dopaminergic cell loss, therefore is not sufficient for assessing complex motor impairment (Braak et al., 2003), seen in Staircase Impairment, and Tube Asymmetry. On the other hand, the Strategy Chaoticity metric revealed substantial changes in motor organization.

**Figure 7.**
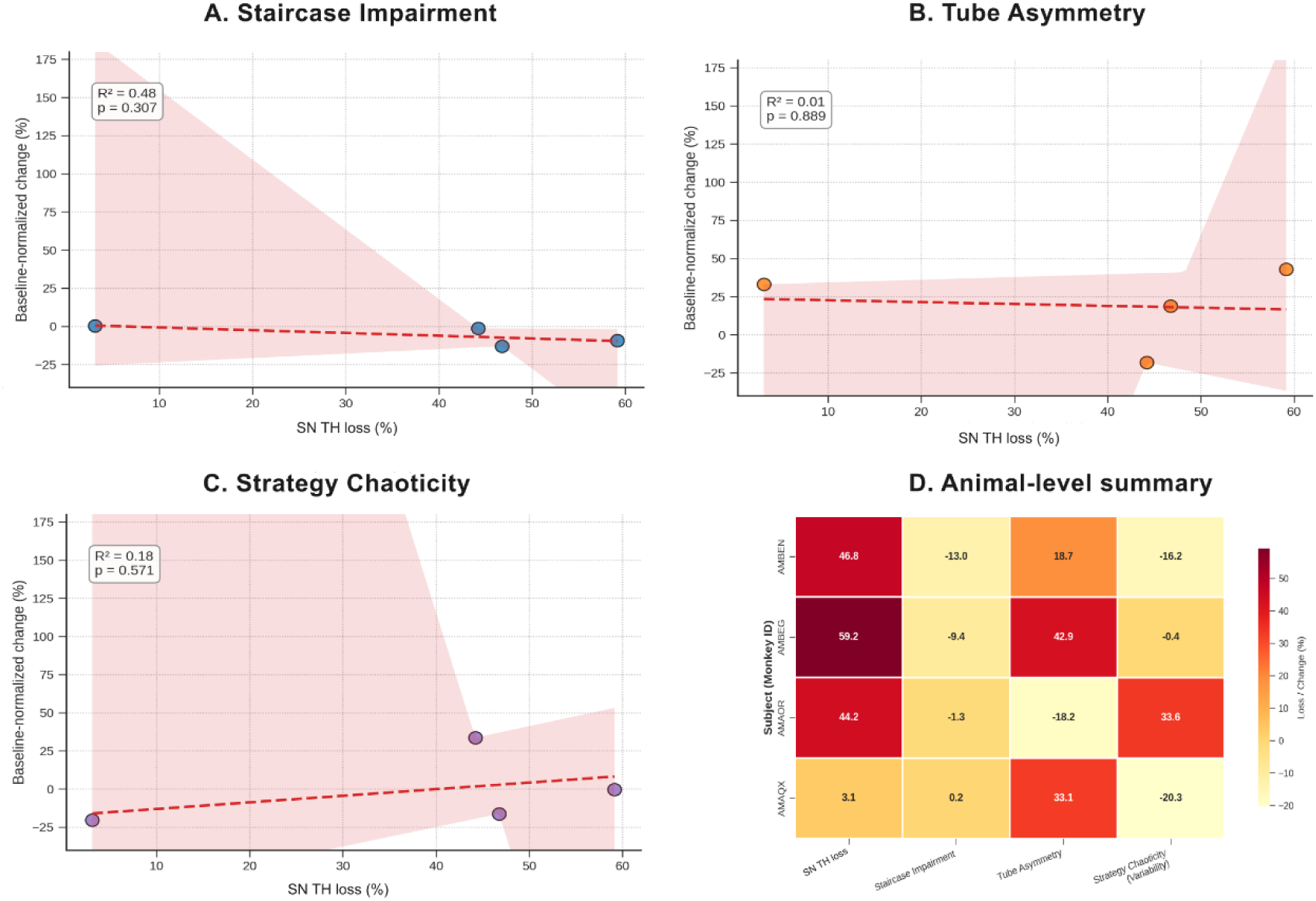
Exploratory association between behavioral impairment and nigrostriatal degeneration. Scatter plots (panels A–C) show the linear regression between SN tyrosine hydroxylase (TH)-positive cell loss (x-axis; %) and baseline-normalized change in behavioral metrics (y-axis; %), individually for each subject. Regression lines (red dashed), 95% confidence intervals (pink shading), R² values, and p-values are displayed within each panel. Given the small sample size (n = 4), analyses are strictly exploratory; effect sizes and directional consistency rather than statistical significance are the primary interpretive focus. (A) Staircase Impairment: baseline-normalized change in reward retrieval success rate on the dominant side (R² = 0.48, p = 0.307). A moderate, non-significant negative trend suggests that greater TH loss is associated with greater staircase impairment, although the relationship is not linear across the concentration range tested. (B) Tube Asymmetry: baseline-normalized change in forelimb use asymmetry between the dominant and non-dominant sides (R² = 0.01, p = 0.889). No meaningful linear association was observed between TH loss and tube asymmetry, consistent with the interpretation that forelimb asymmetry reflects individual compensatory strategies rather than degeneration severity per se. (C) Strategy Chaoticity (Brinkman board deviation variability): baseline-normalized change in session-level deviation variance (R² = 0.18, p = 0.571). The absence of a clear linear relationship is consistent with the heterogeneous nature of motor sequence reorganization across subjects, particularly given BEG HP10’s high pre-lesion baseline variability. (D) Animal-level integrated summary heatmap. Each row represents one subject (AMBEN = BEN HP4; AMBEG = BEG HP10; AOR = AOR HP40; AMAQX = SHAM); each column represents one metric: SN TH Loss (%), Staircase Impairment (baseline-normalized change %), Tube Asymmetry (baseline-normalized change %), and Strategy Chaoticity/Variability (baseline-normalized change %). Color scale reflects the magnitude of each value (darker red = higher absolute value; lighter yellow = values near zero or negative change). Numerical values are displayed within cells. Negative values in behavioral metrics indicate maintained or improved performance relative to baseline, potentially reflecting motor learning, task adaptation, or compensatory dopaminergic upregulation. The heatmap is intended as a qualitative, multidimensional overview to support hypothesis generation rather than statistical inference.

The Animal-level summary can be interpreted as an exploratory multidimensional guide in PD models (Figure 7D). Changes in Tube Asymmetry can be interpreted as improved motor skills (AMAQX-SHAM), or compensatory nigrostriatal degeneration strategy (AMBEN, and AMBEG). It is important to note that sequence-based metrics derived from the Brinkman task showed greater sensitivity to nigrostriatal degeneration.

## 4. Discussion

### 4.1 Proof-of-principle for a precise 6-OHDA model of Parkinsonism in NHP

This study establishes a graded non-human primate model of hemiparkinsonism and demonstrates that motor sequence analysis provides a sensitive readout of dopaminergic dysfunction. There is a need for standardized model systems capable of capturing the heterogeneity of Parkinson’s disease. Current preclinical approaches have relied heavily on rodent models, particularly the 6-OHDA-lesioned rat and the MPTP-treated mouse, as well as transgenic lines overexpressing wild-type or mutant α-synuclein, or carrying loss-of-function mutations in PD-associated genes (LRRK2, PINK1, Parkin, DJ-1) (Emborg, 2007; Zhang et al., 2025). Although rodent models have been instrumental for mechanistic insights, their limited behavioral repertoire constrains the assessment of complex motor functions relevant to Parkinson’s Disease. Several therapeutic candidates for PD —including trophic factors, glutamate receptor modulators, anti-dyskinetic agents, and cell-based therapies—have demonstrated efficacy in rodent models but have shown limited success in Phase II or III clinical trials (Bezard et al., 2025; Vermilyea & Emborg, 2018).

In this context, non-human primates offer a biologically and functionally relevant model, particularly due to the presence of neuromelanin-containing dopaminergic neurons, and a highly developed motor system, thereby more closely reflecting human cellular vulnerability and therapeutic response. In addition, the well-developed prefrontal cortex in primates—disproportionately expanded relative to rodents—supports higher-order cognitive functions, including executive control, working memory, social behavior, and decision-making. This complexity enables NHP models to capture both motor and non-motor dimensions of PD, including cognitive and neuropsychiatric manifestations.

The availability of capuchin monkey (*Sapajus apella*) colonies in Brazil reduces logistical and financial constraints related to breeding and regulatory compliance. The species exhibits a complex motor system, including the ability to use objects from the environment as tools. Consistent with this behavioral repertoire, the motor cortex is highly organized, with well-defined representations of individual digits that support fine motor control. Together, these anatomical and behavioral features enable sophisticated motor performance, as demonstrated in tasks such as the Brinkman board (see Figure 5).

The unilateral model provides the advantage of an internal control, as the contralateral hemisphere serves as a reference within each animal. Based on internal control, TH histological investigation permitted a link between the percentage of neuronal loss and motor behavioral impairments. Performing three different 6-OHDA concentrations (4, 10, and 40 mg/ml), the observed reduction losses were 47%, 59%, and 43%, respectively. Overall, the motor assessment revealed dominant-side motor dysfunction across all tasks, manifested as either increased time to retrieve the reward (Figure 2A) and/or loss of the reward immediately after retrieval (Figure 3A). Importantly, a compensatory strategy was observed in the animals showing the greatest TH reduction (Figure 3D). Furthermore, motor deficits were not fully captured by conventional performance measures, reinforcing the need for more sensitive behavioral metrics.

### 4.2 Behavioral deficits were heterogeneous and captured clinically relevant features of Parkinsonian motor dysfunction

The results demonstrate marked heterogeneity in behavioral deficits following unilateral 6-OHDA induction, despite the use of a standardized surgical protocol. This variability should not be interpreted as experimental noise, but rather as a biologically meaningful feature of the neurobiological response to dopaminergic depletion. In NHP models, the response to partial dopaminergic lesions is well recognized as non-uniform, reflecting inter-individual differences in motor compensatory mechanisms (Emborg, 2007; Teil et al., 2021).

Dopaminergic degeneration does not translate linearly and into motor impairment, as different neuronal subpopulations exhibit selective vulnerability, while pre- and postsynaptic compensatory mechanisms may sustain motor function even in the presence of substantial degeneration (Bezard & Gross, 1998; Brotchie & Fitzer-Attas, 2009; Kish et al., 1988; McGregor & Nelson, 2019; Santana-Román et al., 2025). Nigrostriatal dopaminergic neurons exhibit selective and heterogeneous vulnerability, involving subpopulations with distinct molecular profiles and differential susceptibility to oxidative stress and mitochondrial dysfunction (Blesa & Przedborski, 2014; Lebowitz & Khoshbouei, 2020a). This non-uniform pattern of degeneration directly contributes to the functional variability observed across individuals under similar experimental conditions.

In this context, the observed heterogeneity may be interpreted as the expression of distinct compensatory strategies that vary in efficiency and organization across individuals, as shown in Figures 3C and 3D. Evidence indicates that the motor system can sustain function through pre- and postsynaptic adaptations, including increased efficiency of residual dopaminergic signaling, modulation of neurotransmitter release, and reorganization of basal ganglia circuits(Berardelli et al., 2001; Bezard & Gross, 1998; Brotchie & Fitzer-Attas, 2009; Perez et al., 2008). However, this adaptive capacity is not uniform, resulting in distinct behavioral patterns even under controlled experimental conditions.

Supporting this perspective, both classical and contemporary studies demonstrate that motor symptoms in Parkinson’s disease become clinically apparent only after substantial striatal dopamine loss, typically exceeding 70%, indicating the existence of a functional compensatory phase (Kish et al., 1988; Nutt et al., 2004). Preserved performance in global motor tasks can mask substantial alterations in motor organization and motor strategies, as task execution is maintained through compensatory adjustments that are not captured by conventional performance metrics (Espay et al., 2019).

Preserved task execution does not imply the absence of motor deficits as seen in Supplementary Material 1 but rather reflects behavioral adaptations that partially compensate for dopaminergic dysfunction (Perez et al., 2008; Przedborski, 2017). This finding reinforces the importance of analytical approaches that consider not only final performance outcomes but also movement organization and execution patterns, which are more sensitive to the detection of early motor deficits (Espay et al., 2019).

Taken together, these findings support the notion that inter-individual variability is an inherent feature of hemiparkinsonian models in non-human primates and represents, in fact, a translational advantage. PD itself is characterized by substantial heterogeneity in both dopaminergic loss and clinical manifestation, driven by differences in neuronal vulnerability, molecular factors, and circuit-level reorganization (Blesa & Przedborski, 2014; Santana-Román et al., 2025). Hence, the results suggest that the experimental model employed may capture this variability, representing a promising approach for investigating neural compensation mechanisms and for developing therapeutic strategies targeting not only neuroprotection but also the modulation of adaptive motor processes.

### 4.3 Brinkman board performance provided a sensitive readout of fine motor impairment

The Brinkman board task provided the most sensitive readout of motor impairment in this study, as it constituted a particularly sensitive measure for detecting fine motor deficits, especially because it enabled the analysis not only of overall task outcome but also of spatiotemporal organization of motor execution. Unlike conventional performance measures, based solely on the total number of pellets retrieved, the modified Brinkman board has been widely used as a quantitative tool for assessing manual dexterity, requiring precise grip control, digital coordination, and sequential organization of actions (Savidan et al., 2017; Schmidlin et al., 2011). Furthermore, sequence analysis of pellet retrieval, particularly when represented through heatmaps, enhances the interpretative capacity of the task by revealing disruptions in motor organization even when task success was preserved (Figure 5B).

The Brinkman board allows the dissociation of different components of fine motor control, including precision, coordination, and sequential organization, and is capable of identifying alterations in execution strategy even in the absence of evident quantitative deficits (Badoud et al., 2016; Hoogewoud et al., 2013; Maetzler et al., 2024). In experimental models involving NHP, the modified Brinkman board has been widely employed not only as a quantitative measure of manual dexterity but also as a sensitive tool for investigating functional recovery and motor reorganization following central nervous system lesions(Badoud et al., 2016; Darling et al., 2018; Hoogewoud et al., 2013; Lebowitz & Khoshbouei, 2020b; Nudo, 2013; Savidan et al., 2017; Schmidlin et al., 2011). In this regard, evidence suggests that recovery of performance in manual dexterity tasks does not necessarily correspond to restoration of the original motor pattern but may instead occur through the adoption of alternative compensatory strategies (Hoogewoud et al., 2013; Schmidlin et al., 2011).

A key finding of this study is the dissociation between preserved task performance and disrupted motor sequence organization. Despite the maintenance of task execution in paradigms such as the Staircase test, detailed analysis of the Brinkman task revealed consistent reorganization of motor sequences, suggesting that compensatory mechanisms sustain performance at the expense of movement efficiency and structural organization. This pattern is consistent with the literature on dopaminergic compensation, in which pre- and postsynaptic adaptations, as well as reorganization of basal ganglia circuits, allow preservation of motor function despite significant neuronal loss (Berardelli et al., 2001; Brotchie & Fitzer-Attas, 2009; McGregor & Nelson, 2019; Perez et al., 2008). Thus, preserved global performance does not imply absence of motor deficit, but rather reflects behavioral adaptations that mask underlying alterations in movement organization.

From a translational perspective, findings related to fine motor coordination are particularly relevant, as accumulating evidence indicates that subtle motor alterations emerge in very early stages of PD, even before clinical diagnosis. Studies have shown that fine motor deficits can be detected years before the onset of classic symptoms, particularly involving alterations in finger tapping, handwriting, and manual coordination (Panyakaew et al., 2023; Schaeffer et al., 2024). Consistently, high-resolution digital approaches, such as touchscreen typing analysis, have demonstrated the ability to detect subtle motor impairments associated with Parkinson’s disease, even in early or at-risk stages, revealing alterations in movement kinematics and variability (Iakovakis et al., 2020). Furthermore, deficits in fine motor organization and movement sequencing are already present in the prodromal stage and can be detected through sensitive quantitative metrics even in the absence of overt functional impairment (Yang et al., 2025).

Therefore, the ability of this task to detect alterations in motor sequence organization, even in the absence of deficits in global performance, closely parallels the pattern observed in early stages of Parkinson’s disease in humans, in which task execution is preserved but movement kinematics, timing, and efficiency are already impaired (Berardelli et al., 2001; Espay et al., 2019).Taken together, these findings reinforce the value of the Brinkman board as a sensitive tool for detecting nigrostriatal dysfunction in intermediate or subclinical stages of dopaminergic degeneration, contributing to the development of experimental approaches with greater translational relevance.

### 4.4 Study limitations and model constraints

Parkinson’s disease is characterized by both the progressive degeneration of dopaminergic neurons in the SN and the widespread distribution of Lewy pathology, reinforcing the need for models capable of capturing this complexity (Herculano-Houzel, 2009). Neurotoxin-based models of Parkinsonism, including 6-OHDA, do not fully recapitulate α-synuclein pathology or the progressive nature of the disease. Still, the 6-OHDA model in NHPs induces retrograde dopaminergic depletion, which may offer a valuable window for investigating mechanisms of neurodegeneration and testing neuroprotective strategies (Eslamboli et al., 2005). These limitations should be considered when interpreting the translational scope of the model.

Since 6-OHDA does not cross the blood-brain barrier, the model depends on direct intracerebral delivery. Even minimal surgical deviations may affect the final distribution of the toxin within the target region, thereby contributing to variability in its spread, neuronal uptake, and ultimately the extent of dopaminergic depletion. MRI-based targeting is essential to improve the accuracy of injection placement, particularly in non-human primates, where inter-individual neuroanatomical variation may limit the precision of stereotaxic coordinates derived solely from atlases (Pedrosa et al., 2024). As a proof-of-principle, the present study nonetheless provides a reliable foundation for future methodological refinement, supporting the optimization of targeting precision, lesion consistency, and reproducibility in larger cohorts. Together, our findings highlight the importance of integrating multidimensional behavioral metrics in preclinical models of Parkinson’s disease.

### 4.5 Future directions

From a broader translational perspective, the findings align with proof-of-principle evidence supporting the need for preclinical models capable of capturing the complexity and heterogeneity of PD beyond dopaminergic loss alone. Disease-related dysfunction involves subtle and progressive alterations in motor control, circuit-level reorganization, and integration between motor and cognitive domains, features that are not adequately reproduced by conventional rodent models (Bezard et al., 2025; Emborg, 2007). In this context, recent position statements, including those from the PD-AGE task force, reinforce that NHP models occupy a unique and indispensable role in translational neuroscience, as they preserve key anatomical and functional characteristics of cortico-basal ganglia circuits, enabling the investigation of fine motor control, behavioral adaptation, and progressive dysfunction with greater fidelity to the human condition (Bezard et al., 2025; Teil et al., 2021). Furthermore, the extended lifespan of NHP and their ability to model aging-related processes provide an additional advantage for studying prodromal stages of neurodegeneration and disease progression (Colman, 2018; Mattison & Vaughan, 2017).

## 5. Conclusion

The present proof-of-principle study demonstrates that unilateral stereotaxic injection of 6-OHDA into SN of adult Sapajus apella produces a graded and histologically validated HP model, with ipsilateral tyrosine hydroxylase-positive neuron losses ranging from 44 to 59% relative to the intact contralateral hemisphere, and no meaningful hemispheric difference in the vehicle-injected control animal. This within-subject design, enabled by the unilateral lesion paradigm, provided a robust internal reference for both behavioral and histological comparisons.

Critically, the behavioral findings reveal a dissociation between preserved global task performance and disrupted motor sequence organization: while gross outcome measures such as task completion rates remained largely intact, analysis of fine motor sequence organization using a deviation variance metric applied to the Brinkman board identified post-lesion increases in retrieval sequence disorganization, demonstrating that conventional behavioral measures may underestimate underlying motor dysfunction. This pattern, consistent with dopaminergic compensation mechanisms previously described in both clinical and experimental contexts, emphasizes the inadequacy of task-success metrics as sole endpoints in preclinical models of Parkinson’s disease and highlights motor sequence analysis as a sensitive biomarker of subclinical nigrostriatal dysfunction.

Together, these findings support the translational utility of capuchin monkeys as a biologically relevant NHP model for Parkinson’s disease research and establish a methodological framework integrating quantitative stereology with multidimensional behavioral analysis, as well as highlight motor sequence analysis as a promising biomarker for detecting subclinical nigrostriatal dysfunction. Future studies employing larger cohorts, extended post-lesion timelines, and expanded non-motor assessments will be essential to consolidate these proof-of-principle observations and to evaluate the responsiveness of the motor sequence biomarker to pharmacological or neuroprotective interventions.

## 6. Conflict of interest

The authors declare that the research was conducted in the absence of any commercial or financial relationships that could be construed as a potential conflict of interest.

## 7. Author contributions

(1) Research project: A. Conception. B. Organization and execution.

(2) Data analysis: A. Design. B. Refinement and statistical analysis. C. Review.

(3) Manuscript: A. Writing. B. Review and Critique.

L.R.R.P.: 1B, 2A, 2B, 2C, 3A, 3B.

L.C.P.L: 1B, 2A, 2B, 2C, 3A, 3B.

J.A.P.C.M: 1B, 2C, 3C.

A.G.S: 2B, 2C.

R.R.L: 1B, 3B.

D.SM: 1B, 3B.

B.D.G: 1B, 2C, 3B.

L.V.K: 1A, 1B, 2A, 3A, 3B.

## 8. Acknowledgments

We would like to express our gratitude to the staff of the Medical Clinic in Castanhal-PA. We extend our thanks to the technicians of the National Center of Primates in Ananindeua-PA for their skillful assistance with MRI imaging acquisition, and to collaborators of the Eduardo Oswaldo-Cruz Neurophysiology Laboratory of the Federal University of Pará (UFPA). We are also deeply grateful to Professor Cristovam Wanderley Picanço Diniz and the team of the Laboratory of Investigations in Neurodegeneration and Infection (LNI), Institute of Biological Sciences, Federal University of Pará, for generously providing access to their microscopy infrastructure and for their expert technical support during the histological imaging procedures.

## 9. Data Availability Statement

The datasets supporting the findings of this study are available as supplementary files accompanying the manuscript. These include: raw and session-level Brinkman board performance metrics (brinkman_session_metrics.xlsx), deviation variance summary statistics for the Brinkman board sequence analysis (brinkman_deviation_variance_summary.csv), Wilcoxon test results for Brinkman board sequence variability comparisons (brinkman_variability_wilcoxon_summary.xlsx), Brinkman board retrieval sequence matrices used to generate heatmap visualizations (brinkman_heatmap_matrices.xlsx), Wilcoxon test results for Staircase test parameters (staircase_wilcoxon_results.csv), and Wilcoxon test results for Tube test parameters (tube_wilcoxon_summary.csv). The custom analysis algorithms and scripts used for sequence deviation scoring and statistical processing are not publicly deposited but are available upon reasonable request to the corresponding author, Bruno Duarte Gomes, Laboratório de Neurofisiologia Eduardo Oswaldo Cruz, Instituto de Ciências Biológicas, Universidade Federal do Pará, Avenida Perimetral, 2-224, Room 238, Guamá, Belém – PA, Brazil, 66077-830.

## Notes

### Competing Interest Statement

The authors have declared no competing interest.

### Funding

- This work was supported by the STARDUST project (Grant Agreement No. 767092), funded under the European Union’s Horizon 2020 Research and Innovation Programme, and by a parliamentary amendment from the Government of the State of Pará, Brazil, awarded by State Deputy Dr. Wanderlan Quaresma. The STARDUST project aims to develop a miniaturized, wireless implantable device capable of integrating optogenetic stimulation, electrophysiological recording, and local drug delivery for the modulation of neural circuits associated with Parkinson’s disease. At the Universidade Federal do Pará (UFPA), Brazil, the research component reported in this study — comprising the development and validation of a non-human primate model of experimental parkinsonism for translational application of the device and functional evaluation of the implicated neural circuits — was led by Lane V. Krejcová (Laboratório de Neurofisiologia Eduardo Oswaldo Cruz, Instituto de Ciências Biológicas, UFPA, Belém, Brazil).

