## Supplementary_Figure for "Motor sequence analysis as a sensitive biomarker of dopaminergic degeneration in a non-human primate model of parkinsonism"

**A**

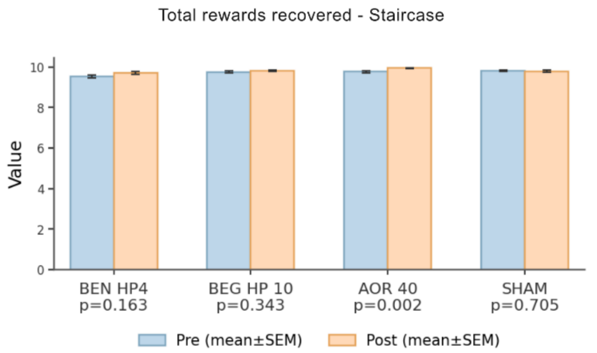


**B**

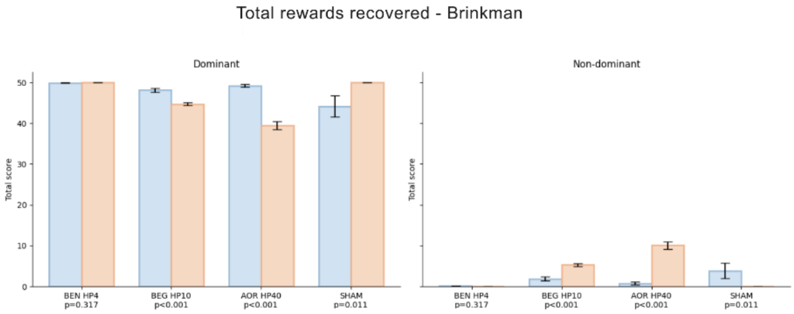


**Supplementary figure.** Conventional performance-based metrics reveal limited sensitivity to nigrostriatal degeneration across the motor task battery. (A) Staircase test: overall reward retrieval success rate. Bar graphs show the proportion of successfully retrieved rewards per session on the dominant (left panels) and non-dominant (right panels) sides before (blue) and after (orange) surgical intervention. Each subplot corresponds to one subject: BEN HP4 (6-OHDA, 4 mg/mL), BEG HP10 (6-OHDA, 10 mg/mL), AOR HP40 (6-OHDA, 40 mg/mL), and SHAM (vehicle-injected control). Data are presented as mean ± SEM. P-values from Wilcoxon signed-rank tests (pre vs. post within-subject comparisons) are displayed below each subject label. Despite confirmed ipsilateral dopaminergic cell losses ranging from 44 to 59% in 6-OHDA-treated animals, overall task completion remained largely preserved across subjects and sides, illustrating the insensitivity of aggregate success rate to underlying nigrostriatal dysfunction. N = 120 sessions per phase (Staircase). (B) Brinkman board: total pellet retrieval score. Bar graphs show the total number of pellets retrieved per session using the dominant (left panels) and non-dominant (right panels) hands before (blue) and after (orange) surgical intervention, for each subject. Data are presented as mean ± SEM. P-values from Wilcoxon signed-rank tests (pre vs. post within-subject) are shown below each subject label. Although some within-subject differences in total score reached statistical significance in BEG HP10 and AOR HP40, the overall magnitude of change was modest and not systematically related to lesion severity. These results contrast with the marked post-lesion disruption in motor sequence organization detected by the deviation variance metric (see Figure 5), underscoring that task completion rate is insufficient as a sole endpoint for capturing the full extent of motor dysfunction in hemiparkinsonian non-human primates. N = 25 sessions per phase (Brinkman board). HP, hemiparkinsonism; SEM, standard error of the mean; 6-OHDA, 6-hydroxydopamine.
